# Amplitude modulation pattern of rat distress vocalisations during fear conditioning

**DOI:** 10.1101/2023.01.18.524509

**Authors:** Eugenia Gonzalez-Palomares, Julie Boulanger-Bertolus, Maryne Dupin, Anne-Marie Mouly, Julio C. Hechavarria

**Affiliations:** Institute for Cell Biology and Neuroscience, Goethe University, 60438 Frankfurt am Main, Germany; Université Claude Bernard Lyon 1, CNRS, INSERM, Centre de Recherche en Neurosciences de Lyon CRNL U1028 UMR5292, CMO, F-69500, Bron, France

## Abstract

In humans, screams have strong amplitude modulations (AM) at 30 to 150 Hz. These AM correspond to the acoustic correlate of perceptual roughness. In bats, distress calls can carry AMs, which elicit heart rate increases in playback experiments. Whether amplitude modulation occurs in fearful vocalisations of other animal species beyond humans and bats remains unknown. Here we analysed the AM pattern of rats’ 22-kHz ultrasonic vocalisations emitted in a fear conditioning task. We found that the number of vocalisations decreases during the presentation of conditioned stimuli. We also observed that AMs do occur in rat 22-kHz vocalisations. AMs are stronger during the presentation of conditioned stimuli, and during escape behaviour compared to freezing. Our results suggest that the presence of AMs in vocalisations emitted could reflect the animal’s internal state of fear related to avoidance behaviour.

## Introduction

Juvenile and adult rats emit two basic types of ultrasonic vocalisations (USVs), with carrier frequencies around 22 and 50 kHz, expressing negative and positive emotional arousal, respectively^1,2^. 50-kHz USVs are emitted in positive interactions, such as playing^3,4^, mating^5,6^ or as response to rewarding pharmacological compounds^7^. On the other hand, 22-kHz USVs are emitted in aversive situations, such as confrontation with predators^8^, submission in inter-male fighting^9^, social isolation^10^, and aversive stimuli^11–13^, including unconditioned and conditioned stimuli. These USVs serve as “alarm calls” to other conspecifics^14,15^ and are part of the rats’ defensive behaviour repertoire.

Though rat vocalisations have been investigated extensively, at present, it remains unknown whether they carry amplitude modulations (AMs). AMs occur in fearful vocalisations of other species, such as human screams^16^ and bat distress calls^17,18^. One hypothesis that has been put forward in recent years is that AMs represent an acoustic correlate of perceptual roughness across species^16,17,19^. Based on this notion, it could be postulated that the 22-kHz USVs of rats, which are typically emitted in aversive situations, also contain AMs.

To test this idea, in this article, we searched for AMs in an existing dataset of USVs emitted during a Pavlovian fear conditioning experiment^20,21^, in which an aversive unconditioned stimulus (US = foot-shock) was presented following a neutral conditioned stimulus (CS = peppermint odour cue). Fear conditioning is widely used in studies of memory and associative learning^22–25^. The strength of the fear memory is commonly assessed using overt behaviour analysis, such as freezing responses. We hypothesised that the AM pattern of 22-kHz USVs could provide additional information related to the animal’s fear response. It is known that other acoustic properties such the duration of USVs and the rate at which these calls are produced correlates well with the freezing rate^13,26,27^. Additionally, these parameters follow a dose-response curve, i.e., the call rate, total call duration and call amplitude increase with increasing shock intensities, while latency of the first call and inter-call length decrease^13,27^.

We report that, as hypothesised, rat 22-kHz USVs have amplitude modulations at rates between 20-100 Hz. In rats, the AM power is highest in vocalisations uttered during the conditioned stimulus odour presentation and in late vs. early trials of the fear conditioning paradigm. We argue that the AM pattern of USVs can be used to obtain information on the rats’ fear response.

## Results

### General characteristics of vocalisations emitted during fear conditioning

The data analysed here have been evaluated on different parameters in previously published studies^20,21^. Note that in the former articles this dataset was not used to search for AMs in the vocalisations, which is the main focus of this paper.

We analysed the vocalisations emitted by rats while undergoing an odour fear conditioning paradigm (Fig. 1A). The trials started with a 30-s long pre-odour period, followed by the odour presentation (CS = peppermint odour which was initially neutral). There were two groups of rats, the first one received 20 s of odour presentation (n = 11), and the second 30 s (n = 9). The presence of two experimental groups is due to methodological differences in the experimental paradigms used in the original publications that collected the data^20,21^. Since these differences are only related to CS duration (20 vs 30 s) we decided to pool together the USVs emitted during the odour presentation in both experimental groups, and to overcome the differences in CS duration by randomly sampling from time periods that have different durations when necessary.

**Figure 1.**
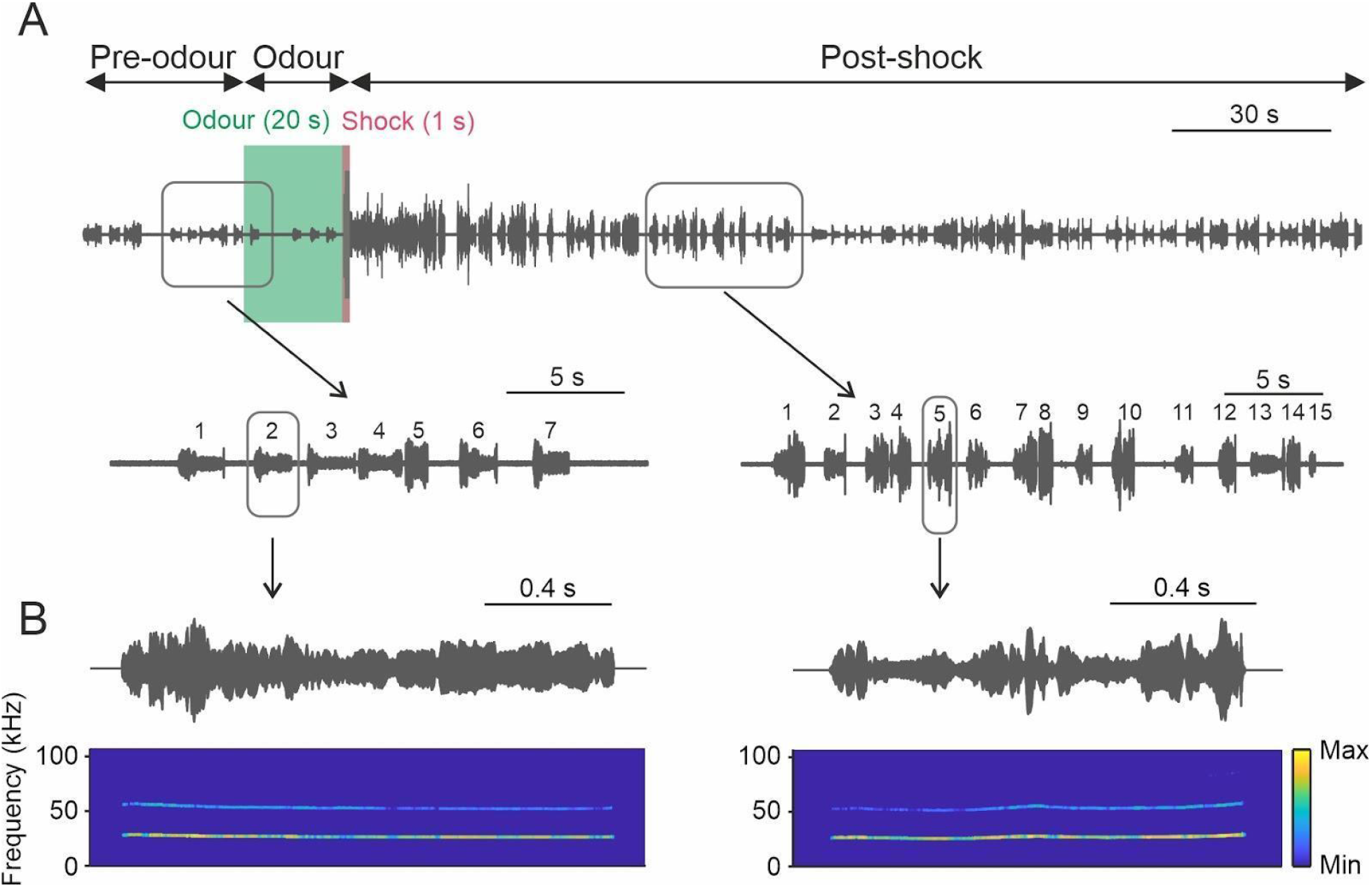
Example of USV recordings during a trial. A) Top: USV were recorded during the 4-min trial divided into three periods: pre-odour, odour, and post-shock. The shock was applied during the last second of odour presentation. Bottom, magnified oscillograms of the recording on the time intervals signalled on the traces above. Each number indicates individual vocalizations. B) Top, oscillograms of two individual vocalisations (zoom-in of A). Bottom, spectrograms of the two vocalisations.

During the last second of odour presentation the rats received a foot shock, that is, the duration of the odour presentation without a shock overlap was 19 and 29 s long for each group, respectively. The following period is referred to as the post-shock period, starting right after the shock, from which we considered the first 30 s for analyses. In total, each trial lasted 4 min. All animals underwent 10 trials, but only the first three and last three trials were analysed. For simplicity, the first and last 3 trials were termed as early and late trials, respectively. We chose to specifically examine early and late trials in order to investigate potential variations that may arise between the initial and later stages of the fear-conditioning task. The vocalisations analysed were at least 300-ms long and their peak frequency was between 18 and 32 kHz. In total, 3353 vocalisations were analysed. An example of a trial recording is given in Fig. 1A, showing two magnified segments with 7 and 15 vocalisations, respectively, and examples of two individual vocalisations in Fig. 1B.

Several parameters of the 22-kHz calls were analysed over the time course of the trials. The idea was to identify the influence of the pre-odour, odour, and post-shock periods on the acoustic parameters of the calls emitted by the rats. Because USV emission strongly impacts respiration^26–29^, the respiration rate was also analysed. The first three and last three conditioning trials from both groups of rats were merged together (group 1: 20 s of odour presentation (n = 11), group 2: 30 s odour presentation (n = 9)).

The first striking result we observed was that the number of vocalisations changed dramatically between pre-odour, odour, and post-shock periods (Fig. 2A, only considered 20 randomly selected 1-s bins in each group and period, Chi-squared test: *p* = 3.5·10^−30^; see Methods for more details). The rats emitted the fewest vocalizations during the presentation of the odor. (Bonferroni-corrected chi-squared tests: *p* = 9.2·10^−20^; *p* = 1.0·10^−30^, compared to pre-odour and post-shock, respectively). Rats vocalized the most during the post-shock period (*p* = 0.04, compared to pre-odour). The duration of the vocalizations during the odour presentation was significantly longer than in the pre-odour and post-shock periods (Fig. 2B; Kruskal-Wallis test: *p* = 6.7·10^−14^, Bonferroni-corrected multiple comparison test; *p* = 5.0·10^−10^; *p* = 1.5·10^−13^, respectively; pre-odour vs post-shock, *p* = 0.19). The vocalizations during odour have the smallest root mean square amplitude (RMS, relative measure of loudness; Fig. 2C; Kruskal-Wallis test: *p* = 2.2·10^−23^, Bonferroni-corrected multiple comparison test; *p* = 2.2·10^−5^; *p* = 1.3·10^−22^, vs. pre-odour and post-shock). Post-shock USVs had the largest RMS (*p* = 1.1·10^−11^, vs pre-odour). The peak frequency was not significantly different between the periods studied (Fig. 2D; Kruskal-Wallis test: *p* = 0.07). Regarding the respiration rate, it was highest during the post-shock period (Fig. 2E; Kruskal-Wallis test: *p* = 1.8·10^−16^, Bonferroni-corrected multiple comparison test; *p* = 6.1·10^−17^; *p* = 4.0·10^−5^, vs pre-odour and odour, respectively), followed by the odour period (*p* = 0.015, vs pre-odour).

**Figure 2.**
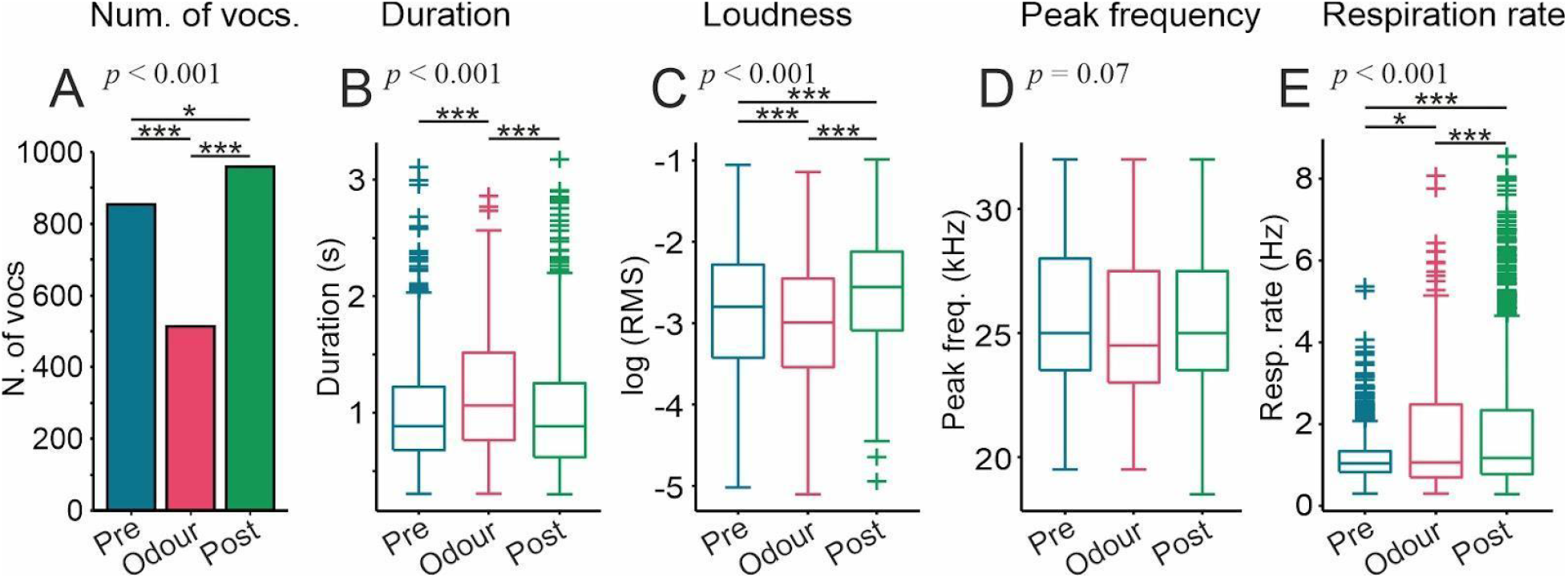
Properties of 22-kHz USVs and respiration rate during the fear-conditioning task. A) Histogram of number of 22-kHz USVs uttered in each period (pre-odour, odour, and post-shock). Chi-squared tests with number of vocalisations in 20 randomly selected 1-s bins. B-D) Boxplots of the call duration, root mean square (RMS) and peak frequency of the same vocalisations. E) Boxplot of respiration rate calculated during 1-s bins. Boxes depict the interquartile range and median. Kruskal-Wallis tests for data shown in B-E (*p* values shown on top of each panel; n = 1300, n = 606, n = 1447, for pre-odour, odour and post-shock, respectively). Stars and horizontal black lines indicate between group comparisons (Bonferroni-corrected rank-sum tests, \**p* < 0.05, *** *p* < 0.001).

We also studied vocalisation onsets for each trial and rat (Fig. S1), to better assess the temporal pattern of USV production. Despite the difference in the number of vocalisations uttered by each individual, a general call rate reduction during odour presentation was noticeable. We also observed a greater call rate in late trials and that the increase in rate in the post-shock period is not restricted to a specific window but is rather a consistent increase throughout the period analysed. The median inter-call interval appeared to be stable across individuals (Fig. S2), despite inter-individual differences in the underlying distributions.

Since the odour period includes one second during which the shock is presented, we performed the same analyses displaying those USVs uttered during the shock and those that were not (Fig. S3A-D). Because of the low number of USVs during the shock, n = 8, statistical tests were not performed. However, it is evident that both shock-related and odour-related vocalisations fall within a similar range (for the variables measured) when examined qualitatively.

Recently a new type of aversive vocalisation has been described in rats, the 44-kHz vocalisations^30^, which are long (> 150 ms) and with peak frequencies > 32 kHz. We searched for vocalisations that met these criteria (> 150 ms; 35-72 kHz) and found 43 calls (Fig. S4A-C). Their peak frequency was 36.6 ± 1.9 kHz (mean ± std) and they were uttered predominantly in late trials and during the post-shock period.

### Changes in acoustic parameters across trials

In order to investigate the effect of previous exposures to the CS-US pairs, the acoustic parameters mentioned in the preceding text were analysed in a trial by trial manner (considering trials 1 to 3 and 8 to10; Fig. 3). This analysis was done for each trial period separately (i.e., pre-odour, odour, and post-shock). Again, here both experimental groups (20 and 30 s odour presentation) were merged together. We observed the following:

**Figure 3.**
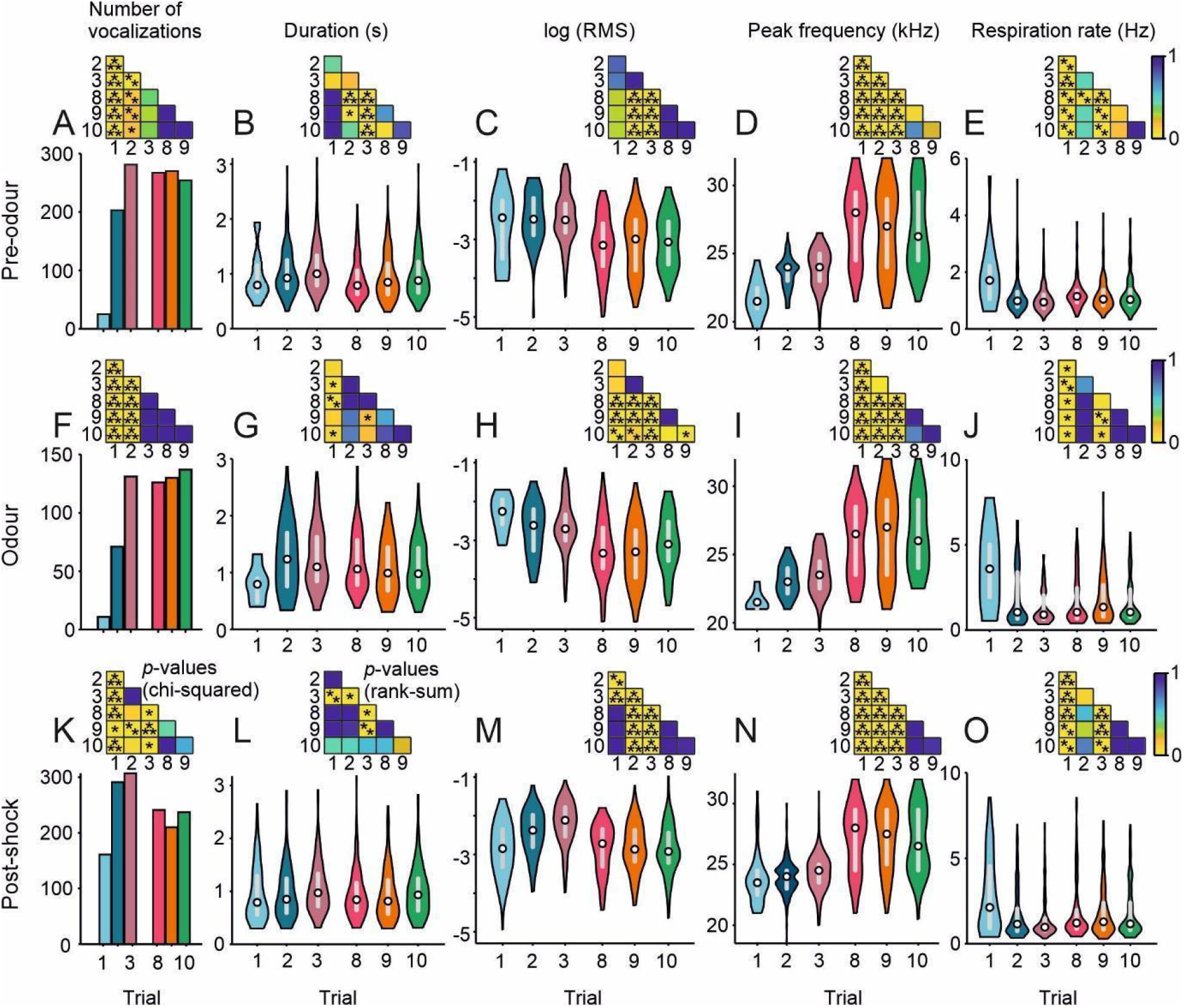
Analysis of 22-kHz USVs as a function of trial number. Histogram of number of vocalisations (A, F, K) and violin plots of duration (B, G, L), root-mean square (RMS; C, H, M), peak frequency (D, I, N) and respiration rate (E, J, O). White circle indicates the median and grey bar the interquartile range. Insets: statistical significance; number of vocalisations: Holm-Bonferroni corrected chi-squared test; duration, RMS, peak frequency, and respiration rate: Holm-Bonferroni corrected Wilcoxon rank-sum tests. * *p* < 0.05, ** *p* < 0.01, *** *p* < 0.001.

1. Number of vocalisations (Fig. 3A, F, K): The number of vocalisations increased in both pre-odour and odour periods in the first two trials reaching a stable level in the third trial, which was maintained in later trials (Chi-square tests, first and second trials compared to the rest, *p* < 0.05). In the post-shock period, a stable number of vocalisations is reached in the second trial.
2. Duration (Fig. 3B, G, L): The duration of the 22-kHz USVs in the pre-odour and post-shock periods was relatively constant. During the odour presentation, the vocalisations were slightly shorter in the first trial than in the rest (Holm-Bonferroni corrected Wilcoxon rank-sum tests, *p* < 0.05).
3. Loudness (Fig. 3C, H, M): The vocalisations were mainly louder (measured as uncalibrated RMS) in the first trials compared to the last ones before and during the odour presentation. In the post-shock period, the vocalisations were loudest in the second and third trials, reaching in later trials comparable levels to the first trial.
4. Peak frequency (Fig. 3D, I, N): The peak frequency increased gradually and significantly in the first three trials in all periods relative to odour presentation. Peak frequency was even higher in the last trials, where it remained more stable.
5. Respiration rate (Fig. 3E, J, O): The respiration rate is highest in the first trial in all three periods reaching the lowest level in the third trial. The later trials have comparable levels to the second trial. These data point towards an inverse relationship between respiration rate and number of vocalisations.

### 22-kHz USVs have fast amplitude modulations

In the following, we describe the analysis pertaining to the amplitude modulation pattern of rat 22-kHz vocalisations, which was at the centre of our interest. We wanted to determine if the rat 22-kHz USVs carry AMs as described in previous studies for bats and humans. We reasoned that calls emitted in different trials and/or trial periods (pre, post and odour presentation) could have different AM patterns.

The method used to evaluate amplitude modulations was the modulation power spectrum (MPS). The same method has been used to study AMs in humans, bats and birds^16,17,19,31^. The MPS allows the analysis of both temporal and spectral modulations simultaneously. In the MPS, positive and negative temporal modulations display downward and upward frequency modulations, respectively.

For the initial MPS analysis, we hand-picked 50 example rat vocalisations across trials and trial periods: 25 vocalisations that displayed strong AM in their oscillogram, and 25 that did not (see examples in Fig. 4A). The objective was to compute the MPS of the example calls, and to compare fast AM vs. slow AM to identify regions of interest (ROIs) in the MPS. The ROIs would indicate MPS regions where slow and fast modulated vocalisations differ from each other. We thought that once identified, these ROIs could be used in all subsequent analysis. At the methodological level, for computing the MPS we focussed on the first 300 ms from all vocalisations (RMS-normalised). This approach let us equalise the spectro-temporal resolution of the analysis across vocalisations, and to focus on AM differences rather than on RMS differences.

**Figure 4.**
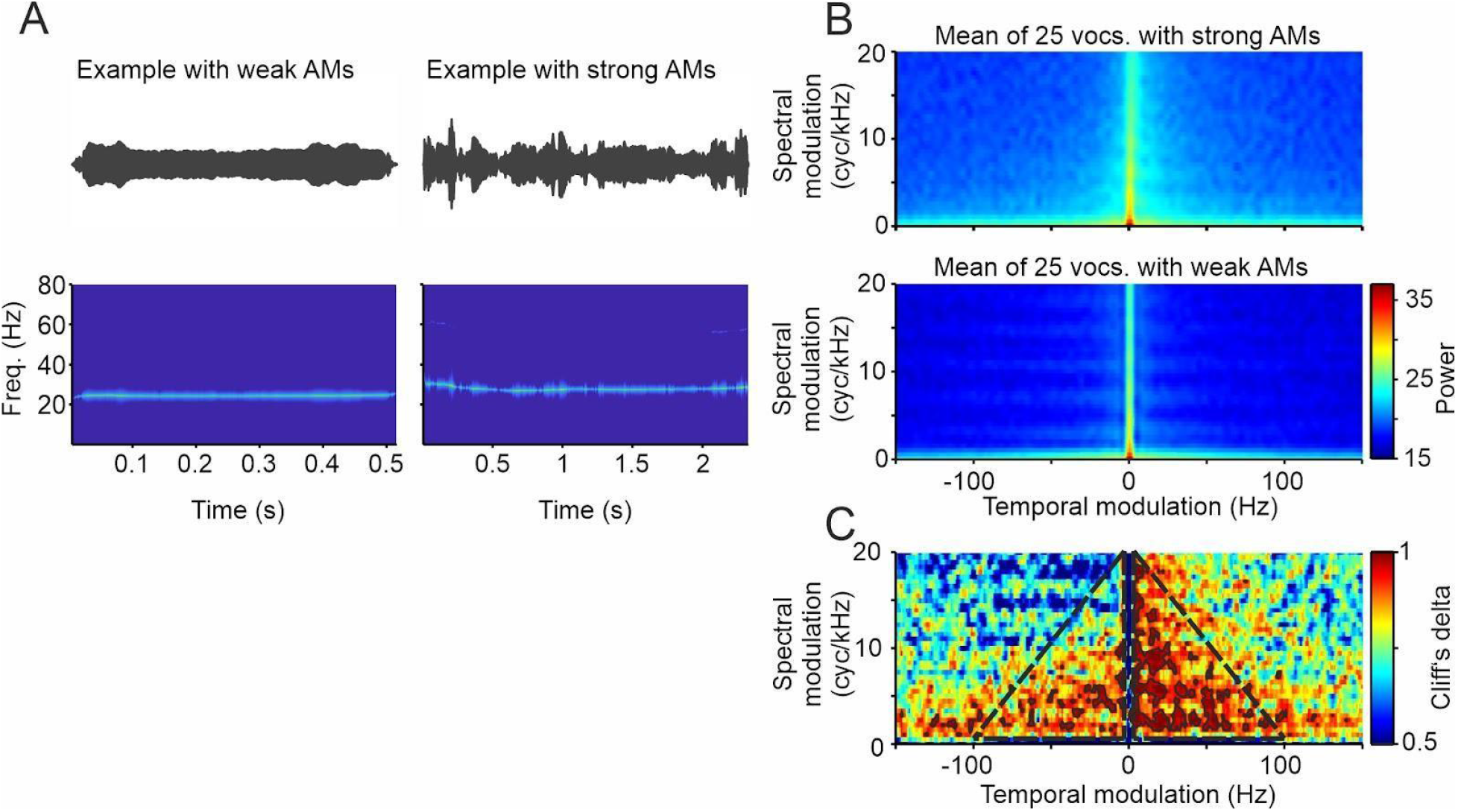
A) Oscillograms (top) and spectrograms (bottom) of two examples of 22-kHz rats’ vocalisations, one with weak (left) and one with strong (right) amplitude modulations (AMs), . B) Mean of the modulation power spectra (MPS) of 25 manually selected vocalisations with strong (A, top) and weak (A, bottom) AMs. Only the first 300 ms of each vocalisation were considered and all were RMS-normalised. C) Cliff’s delta of the MPS of the vocalisations used in B) comparing strong and weak AMs. Solid black lines designating areas with values > 0.948 (double the threshold of what is considered as large effect size). Dashed black lines show the area within which the mean of the MPS values were calculated for all vocalisations (what is considered as AM scores in this manuscript).

Figure 4B shows the average MPS of the 25 hand-selected fast AM rat calls and the 25 slow AM calls (top and bottom, respectively). As it can be noted, the MPS of the fast AM calls is broader than that of the slow AM vocalisations. For comparison purposes, we calculated the Cliff’s delta metric, an effect size measure for group comparisons which has values between -1 and 1, the closer to zero, the more similar the groups^32^. In the resulting Cliff’s delta matrix, we selected the regions that were higher than 0.948: double the absolute value that is typically used to define a large effect size^32^. The more red the values are, the more often those values are larger in the vocalisations with strong AMs. These regions indicate very high differences between the MPS of fast and slow AM calls and are indicated with solid black lines in Fig. 4C. Differences with very large effect sizes occurred consistently between the two groups at frequencies between 3 and 100 Hz, mostly on the right side of the MPS that corresponds to downward frequency modulations. Based on the results obtained, we outlined ROIs (dashed black lines in Fig. 4C) from which we could calculate an AM score. The AM score was computed for each vocalisation as the mean value of the ROIs defined in Fig 4C (temporal modulations: 3-100 Hz, positive and negative; spectral modulations: 1-20 cyc/kHz). The AM score was calculated in the same manner for all the vocalisations.

When considering all the calls, we observed that the AM score was significantly highest during the odour period (Fig. 5A; Kruskal-Wallis test: *p* = 2.5·10^−6^, Bonferroni-corrected multiple comparison test; pre-odour vs odour, *p* = 2.2·10^−6^; pre-odour vs post-shock, *p* = 0.96; odour vs post-shock, *p* = 6.5·10^−5^). When considering the trials separately, the AM score was highest in most cases in late trials during the pre-odour and odour periods (Fig. 5B; Holm-Bonferroni corrected Wilcoxon rank-sum tests, *p* < 0.05), although during the post-shock period this effect was not clear.

**Figure 5.**
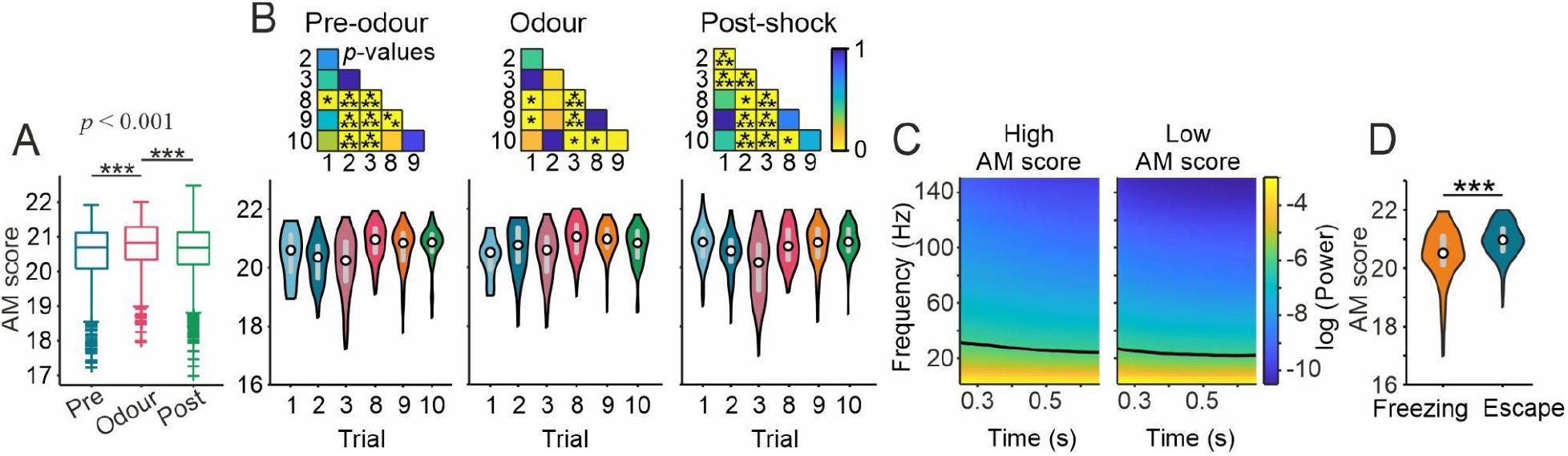
Amplitude modulation score. A) Boxplot of the AM score in each for the three periods during the first three and last three trials for both groups. Boxes depict the interquartile range and the horizontal line inside each box indicates the median. B) Violin plots of the amplitude modulation score (see Methods) for each period and trial. Insets: *p* values of Wilcoxon rank-sum tests (Holm-Bonferroni corrected). White circle indicates the median and grey bar the interquartile range. C) For this panel all vocalisations that are between 0.90 (median) and 1.5 s long were used (1164 in total). The envelope of these vocalisations was calculated, only the first 900 ms was considered and they were z-normalised. Then the spectrograms of the envelopes were calculated. These vocalisations were median split according to their AM score and the mean spectrogram of each group is plotted here. Logarithmic axes. Black lines designate an arbitrary threshold of -6. D) AM score of the vocalisations emitted right after the shock (10 s post-shock period) and classified into freezing (n = 145) or escape (n = 141) depending on the rat’s behaviour (Wilcoxon rank-sum test). * *p* < 0.05, ** *p* < 0.01, *** *p* < 0.001.

We also analysed the mean spectrogram of the envelopes of the vocalisations (Fig. 5C; n=1164; see Methods for more details). This analysis gives access to changes in the AM of individual calls over time. Here we split USVs into two groups, those with weak and strong AMs, respectively, as defined by the AM score of each call and whether it was higher or lower than the population median (median split). We observed that the AMs were higher at the beginning for both vocalisations with strong and weak AMs and remained stable towards the end (Fig. S5).

When the rats received the shock, they displayed escape behaviours rapidly followed by freezing^20^. In order to look for a possible effect of the defence behaviour displayed by the animal (freezing or escape), we compared the AM score between USVs emitted during the 10 s after the shock. We found that the USVs emitted during escape had stronger fast AMs than those emitted during freezing (Fig. 5D; Wilcoxon rank-sum test, *p* = 2.7·10^−10^).

The AM scores of vocalisations uttered during the shock are among the largest values observed (Fig. S3E; no statistics performed due to the low number of shock vocalisations). In a similar way, the AM scores of 44-kHz USVs were also within the high range compared to 22-kHz (Fig. S4D-E; again, no statistics performed, due to low sample size).

The example with AMs (Fig. 4A right) displays also a down frequency modulation at the onset. We found this observation to be consistent across vocalisations (Fig. S6A), and that the downward frequency modulation is correlated with high AM scores (Fig. S6B-C).

### The amplitude modulation score is highest in longer and fainter vocalisations

The acoustic analyses hitherto showed the presence of “fast” amplitude modulations, which are strongest during the odour presentation of fear conditioning trials. To determine how the acoustic parameters may interact between them, we performed correlational analyses considering all 22-kHz USVs measured (Fig. 6; Pearson’s correlation coefficient and linear regression). Considering the strongest correlations, the number of vocalisations was positively correlated with the calls’ peak frequency, while the AM score was negatively correlated with the calls’ loudness (measured as logarithmic RMS; -1 values designate louder calls). The AM score was positively correlated with the respiration rate (Fig. 6F). To further explore this relation, we calculated the correlation between the changes in AM score and respiration rate across periods (Fig. 6G-I). In the post-shock period, the AM score and respiration rate significantly increased compared to the odour period (other period combinations were not significant). For other acoustic parameters, the strength of the correlations was mostly weak as illustrated in Fig. S7 (for the *r*’s interpretation see^33^).

**Figure 6.**
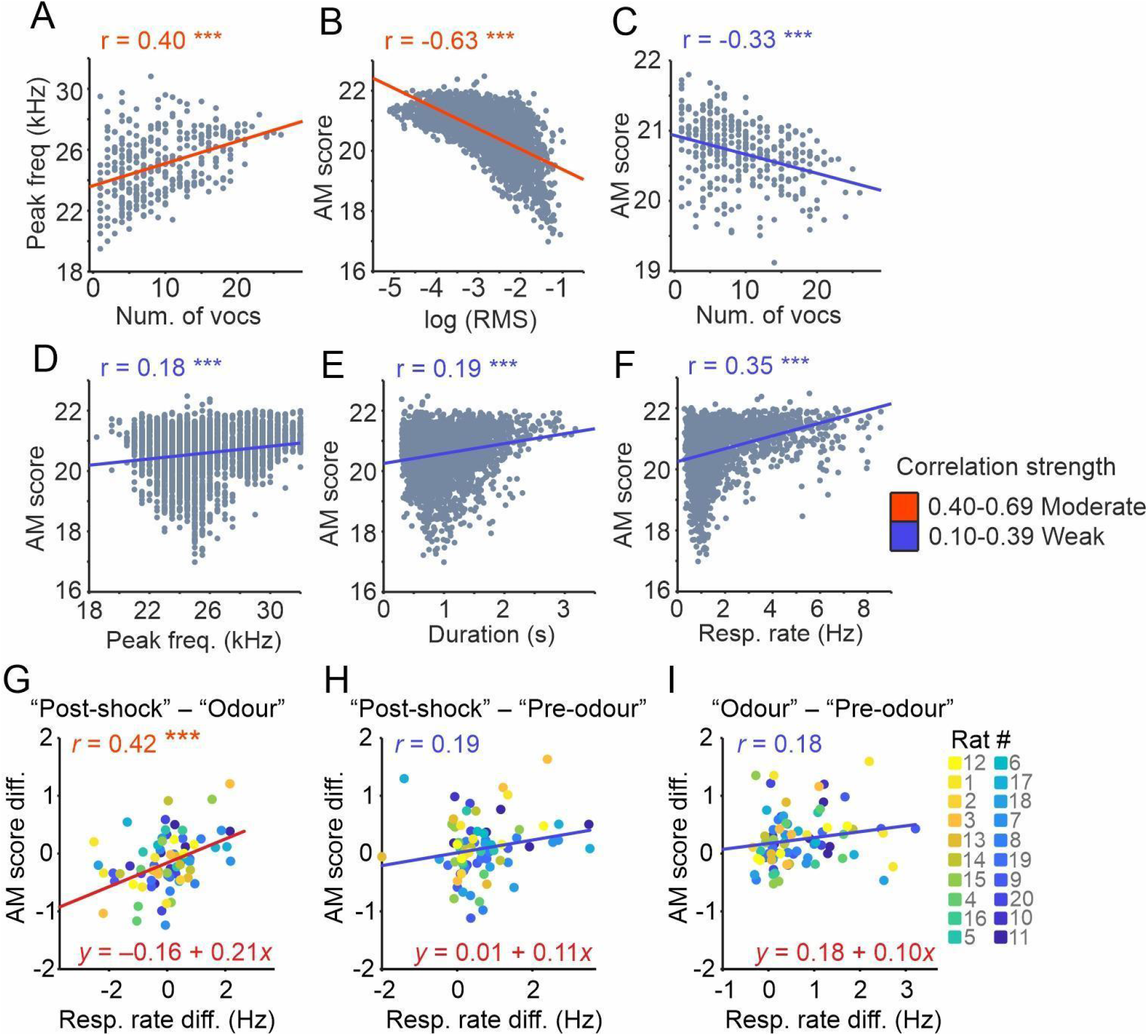
Correlations of 22-kHz vocalisations. A) Peak frequency plotted against number of vocalisations (here it was considered the mean peak frequency and the mean number of USVs emitted during 5-s bins for each rat and trial). B-F) AM score plotted against the natural logarithm of the root-mean square (RMS), number of vocalisations (calculated as the mean in 5-s bins), peak frequency, duration and respiration rate. G-I) AM score difference plotted against the respiration rate difference (“Post-shock” - “Odour”, G; “Post-shock” - “Pre-odour”, H; “Odour” - “Pre-odour”, I). Each dot represents one trial and rat (colour-coded for rat’s identity; see legend). Lines depict the fitted linear regression (see colour legend for the interpretation). Pearson’s correlation coefficient (*r*) shown for each plot. *** *p* < 0.001.

## Discussion

The main goal of this study was to analyse the temporal modulations of rat 22-kHz USVs emitted while undergoing a fear conditioning task, together with the animals’ respiration rate. A specific aim of the study was to investigate whether amplitude modulated sounds (putative roughness) can be detected in rat vocalisations as the level of stress increases in a fear conditioning task.

We found that: (1) during the odour period (compared to the pre-odour period), the number of 22-kHz vocalisations decreases together with the loudness, whereas the calls become longer and the respiration rate increases; (2) during the post-shock period, the number of calls increases (more than in pre-odour) and call duration becomes shorter (as in pre-odour) and the respiration rate increases (more than during odour). (3) In late trials (compared to early), the number of vocalisations increases, the calls’ peak frequency is higher, whereas call loudness is reduced. (4) There are fast amplitude modulations in rats’ 22-kHz USVs. These fast modulations cover the frequency range between 3-100 Hz, and they occur mostly during CS presentation, and generally more during late trials and during escape behaviours.

### Comparison with previous studies

Rats emit most of the 22-kHz USVs when immobile^27^, and call emission occurs together with increases in heart rate and blood pressure^28,34,35^. The majority of the situations in which rats emit 22-kHz USVs have clear aversive components, e.g., confrontation with a predator, during intermale social defeat^36,37^ and during fear conditioning^12,13,38,39^. During the CS presentation (an odour in our case) the call rate decreases, also in line with past literature which shows that USVs are more frequently uttered during the intertrial interval compared to the CS presentation^12,28,40^. It has been suggested that context cues between the CS-US pairings may signal anxiety, whereas the CS prior to the US may indicate acute fear, and that USVs may specifically reflect the anxiety state^12^. While freezing is often similar in fear and anxiety states, sustained 22-kHz USV have been shown to occur preferentially between trials, whereas acute fear induced by the CS as an imminent danger signal resulted in a decrease of the number of USVs^12,28^.

The results presented in this manuscript are consistent with prior findings suggesting suppression of context supported USV by the CS. Additionally, in line with previous studies, we have shown here that the number of vocalisations, as well as the peak frequency, increase with the situation’s aversiveness^41^. Long high-frequency aversive calls, namely 44-kHz^30^, although more uncommon, occur predominantly in late trials, as previously reported^30^, which could be also related to the aversiveness level. Shock delivery restores USV emission levels (to pre-odour levels) and increases the loudness. With increasing number of trials, the number of utterances and their peak frequency increase as well. This is in consonance with previous data showing that call rate increases with the aversiveness level^27,41^.

### Amplitude modulations

A central aim of the present work was to investigate whether amplitude modulations can be found in rat USVs. We brought evidence that such modulations are indeed present especially during odour presentation. According to our results, the amplitude modulation score is negatively correlated with the vocalisation’s loudness and number of vocalisations and positively correlated with the peak frequency, duration and respiration rate. The AM score also increases with the number of trials. According to Morton’s motivational rules, broadband low-frequency sounds are associated more with hostile situations^42^. Since the 22-kHz are very narrow spectrally, the amplitude modulations could be a useful way to increase the information they carry, in a similar way that amplitude modulations generate rough-like sounds in humans^16^. Increasing information content through AMs can be related to richer amplitude changes in the time domain or to frequency side-bands that appear in the vocalisations’ spectra as a consequence of the AMs.

Our study shows that the AM pattern of rats’ 22-kHz USVs can be changed. The AM pattern could reflect the animal’s internal state. Yet, whether rats can willingly control AM presence/absence and whether the AM patterns observed are useful for the listeners remains unclear. Nonetheless, AMs in rat vocalisations could be a useful approach to complement existing metrics that characterise rat behaviour in natural environments and in laboratory studies (i.e., experiments that measure USV calls to study, for example, drug effects).

The defensive responses triggered by the foot shock can be passive (freezing) or active (escape). In a published article using the same data as here^20^, the authors analysed the USVs emitted during the first minute after the foot shock during the two situations (freezing and escape). During escape, the call duration was shorter, and the peak amplitude and frequency were higher than during freezing. Here, we analysed the AM score of the vocalisations uttered 10 s after the shock as well, and found that during escape, the USVs have stronger AMs. These observations suggest that AMs of USVs play a role in the rats’ active defence behaviour. In that sense, AM constitutes a useful bioacoustical marker that could allow to distinguish between USVs signalling acute fear and those reflecting anxiety, and between passive and active defence behaviours.

### Final remarks

Amplitude modulations in human screams and bats distress sounds have been proposed as the correlate of acoustic roughness^16^. Here we showed that AMs (3-100 Hz) also occur in rat 22-kHz aversive calls emitted in a fear conditioning paradigm. AMs in rat USVs are strongest during CS presentation, and AM strength shows a trend to increase after the first shock application, denoting a relation with aversiveness. We noticed that the range of the frequencies of the AM in rats (∼ 3-100 Hz) is not as delimited and high frequency as in bats (AMs centred at 1.7 kHz^17^), but rather covers the same broad range of frequencies that characterises perceptual roughness in humans (∼ 30-100 Hz^16^).

There could be two possible, not mutually exclusive, functions for the AMs found in 22-kHz rat vocalisations. First, they may reflect the internal affective state of the animals such as the fear for a potential threat and/or related to the avoidance behaviour. This is supported by the fact that the AM score increases after the first shock delivery during CS presentations and that it is higher during escape than during freezing after the shock. They may be modulated quantitatively and/or qualitatively by the emotional state (fear and/or avoidance). Second, they might be used as a communication signal to indicate a potential danger to other conspecifics.

## Methods

The data used here were recorded during experiments directed to the investigation of the respiration and brain neural dynamics during odour fear conditioning. Those results are published elsewhere^20,21^. The present article shows analyses of the acoustic recordings, including new insights to amplitude modulations. For a more detailed description of the paradigm, experimental apparatus or data acquisition, please refer to the aforementioned papers^20,21^.

### Animals

The data have been obtained from twenty male Long Evans rats (290-375 g, at the start of the experiments, Janvier Labs). The animals were housed individually at 23 °C, 12 h dark-light cycle, with food and water available *ad libitum*. All procedures were performed in accordance with the European Union guidelines and regulations (Directive 2010/63/EU of the European Parliament and of the Council of the European Union regarding the protection of animals used for scientific purposes), they have been approved by the French Ethical Committee N° 055 and the project is referenced by the French Ministry of Research as APAFIS #10606-2017071309472369v2. The study was conducted in accordance with the ARRIVE (Animal Research: Reporting of *in Vivo* Experiments) guidelines.

### Experimental apparatus

A detailed description of the experimental apparatus can be found here^27^. Briefly, it consisted of a whole-body customised plethysmograph (diameter 20 cm, height 30 cm, emka TECHNOLOGIES, France) inside a sound-attenuating cage (L 60 cm, W 60 cm, H 70 cm, 56 dB background noise). The bottom of the chamber was equipped with a shock floor connected to a programmable Coulbourn shocker (Bilaney Consultants GmbH, Düsseldorf, Germany). On top of the plethysmograph three Tygon tubing were connected to an olfactometer. Deodorized air flowed constantly through the cage (2 L/min). An odour (McCormick Pure Peppermint; 2 L/min; 1:10 peppermint vapour to air) was introduced when scheduled. A condenser ultrasound microphone (Avisoft Bioacoustics CM16/CMPA, Berlin, Germany, sampling frequency 214285 Hz, 16 bit) placed on top of the plethysmograph was used to record the vocalizations, that were analysed offline. Two cameras recorded the animals’ movement. The freezing percentage per 1-s bin was automatically detected from using a Labview software and verified by a researcher.

### Odour fear conditioning

The animals were placed individually in the apparatus for 30 min per day during 3-4 days before the start of the experiments to get familiarised with the device. Before the recording started, the animals were allowed 4 min for exploration, then the odour (McCormick Pure Peppermint; 2 l/min; 1:10 peppermint vapour to air) was applied during 20 or 30 s, the last second of which overlapped with the foot-shock (0.4 mA). The animals underwent a total of 10 trials with an intertrial interval of 4 min. Each trial consists of three periods: 30 s “pre-odour”, 20 or 30 s “odour”, and 190 or 180 s “post-shock”, of which only the first 30 s were considered. Eleven rats always received a 20 s odour delivery and the other nine, 30 s. For the current article, the vocalisations from both groups of animals were analysed together while keeping track of the period in which they were uttered. A total of 6 trials were analysed, early trials (# 1, 2 and 3) and late trials (# 8, 9 and 10). This was resolved to allow us to observe a potential effect of the animal’s expectation in the vocalisation parameters, as the rats become familiar with the paradigm.

### Data analyses

The vocalisations were first detected automatically in Avisoft SAS Lab Pro software (v. 5.2 Avisoft Bioacoustics, Germany) using an amplitude threshold of 1.9 %, minimum duration of 1 ms and hold time of 45 ms. Subsequently, all detections were manually revised and modified when necessary. All further analyses were done in custom-written scripts in Matlab (R2020a, The MathWorks, Massachusetts, USA). Vocalisations that were longer than 300 ms and that had a peak frequency between 18 and 32 kHz (i.e., the so-called 22-kHz vocalisations) were considered for further analyses (3353 from 3988 detected calls). The spectrogram shown in Fig. 1 was calculated using *mtspecgramc* function from Chronux (2 tapers of 50-Hz bandwidth and 0.2-s duration, 5-ms long windows at 4-ms steps). The loudness was calculated as the root-mean square (RMS) and the peak frequency using *mtspectrumc* (2 tapers). The peak frequency in Fig. S6 was calculated with the Chronux function *mtspectrumc* with default parameters.

An amplitude modulation score was calculated from the modulation power spectrum (MPS) of the first 300 ms of all vocalisations. First, the vocalisations RMS normalised. Second, the MPS were calculated as the 2D fast Fourier transform of the spectrograms (Short time Fourier transform, window length = 2048, number of FFT points = 512, hop =1). Subsequently, the MPS were 2D interpolated for averaging purposes (temporal and spectral resolutions: 1 Hz and 0.5 cyc/kHz, respectively, “spline” method). Finally, the score is calculated as the mean power two triangular regions with vertices in (3 Hz, 1 cyc/kHz), (100 Hz, 1 cyc/kHz) and (3 Hz, 20 cyc/kHz) for one triangle, and (-3 Hz, 1 cyc/kHz), (-100 Hz, 1 cyc/kHz) and (-3 Hz, 20 cyc/kHz) for the second one.

For Fig. 5C, were considered those vocalisations with durations between 0.9045 s (median of all vocalisations) and 1.5 s (n = 1164, out of 3353). Only the first 0.9 s were considered. The envelopes of these vocalisations were calculated (secant method, temporal resolution: 0.47 ms), 10 ms rise/fall time and were z-normalised. Then they were downsampled 200 times (sampling frequency: 1071 Hz) and the spectrogram of the envelopes were calculated using the *mtspecgramc* function (5 tapers, 500 ms window, 5 ms step size, no padding, frequency band: 1-150 Hz). The spectrograms were divided into two groups by a median split of the AM score. The figure shows the mean spectrogram of each group with natural logarithmic power. The same analysis was performed for the last 0.9 s of the vocalisations (Fi     g. S5).

The vocalisations that were longer than 150 ms, with a peak frequency between 35 and 72 kHz and after manual revision, were classified as 44-kHz^30^ (Fig. S4; n = 43). The mean spectrogram in Fig. S4A was calculated using *mtspecgramc* (50 ms window, 1 ms step size, default parameters for the rest). For those 44-kHz > 300 ms (n = 37), the AM score was calculated as described above.

In order to compare the score in the two defence behaviours (freezing or escape), USVs emitted during the 10 s after the foot shock were analysed. USVs emitted during freezing were those whose onset was in time bin (1-s long) with ≥ 50 % of freezing, and the rest was considered as escape.

### Statistical analysis

Statistical analyses were done in Matlab. The normality of data distributions was assessed by the Kolmogorov-Smirnoff test. Since none of them were normal, non-parametric tests were used. To compare the number of vocalisations (Fig. 2A) between the different periods (pre-odour, odour, and post-shock), twenty random bins of 1-s duration were selected for each period so that they had the same number of bins for chi-squared tests (significance at *p* < 0.05). *P* value shown is the average after one hundred repetitions. For comparisons between two periods, the same procedure was performed with Holm-Bonferroni-corrected chi-squared tests. Statistical differences for the duration, log (RMS), peak frequency, respiration frequency (Fig. 2B-E) and amplitude modulation score (Fig. 5A), were calculated using Kruskal-Wallis tests (significance at *p* < 0.05). If significant, a multiple comparison test was performed (Bonferroni corrected; *multcompare* in Matlab). For comparisons between different trials, Wilcoxon rank sum tests (significance at *p* < 0.05) were used. For comparisons of values between the same vocalisations (Fig. S6), the signed rank test was used.

## Acknowledgements

We thank our funding sources: Deutsche Forschungsgemeinschaft grant no. (428645493) (to E.G.P and J.C.H); Centre National de la Recherche Scientifique and the LABEX CORTEX (ANR-11-LABX-0042) of Université de Lyon, within the program “Investissements d’Avenir” (ANR-11-IDEX-0007) operated by the French National Research Agency (to J.B.B, M.D. and A.-M.M.).

## Data availability statement

The original data of this study are available from A.-M.M. upon reasonable request.

## Author contributions

M.D., J.B.B. and A.-M.M. conceived the experiments. M.D. carried out the experiments; E.G.P., J.B.B., A.-M.M. and J.C.H. designed the study; E.G.P. analysed the data; E.G.P. wrote the first draft of the manuscript; all authors revised the manuscript.

