## Supplementary Figures for "Amplitude modulation pattern of rat distress vocalisations during fear conditioning"

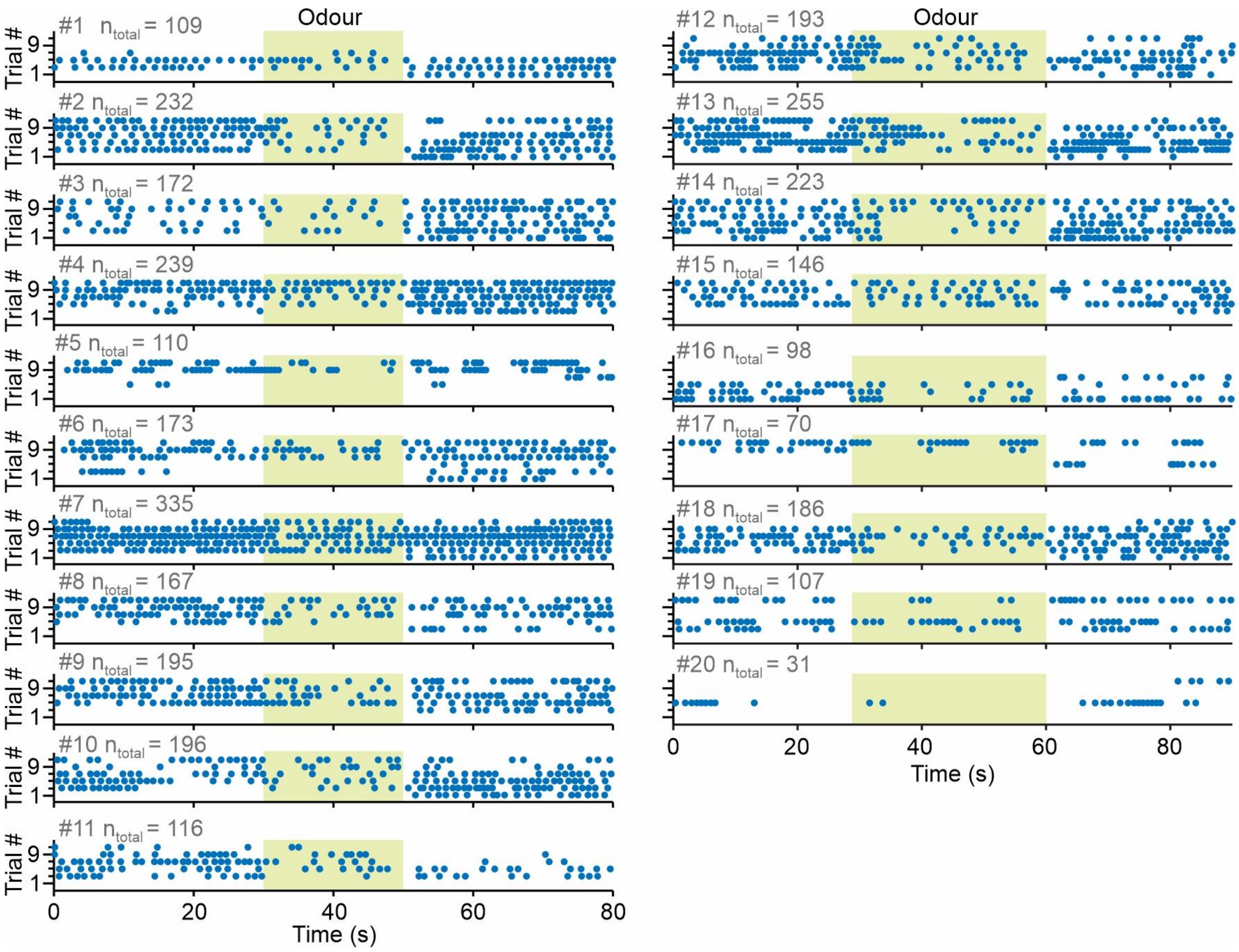


Figure S1. Scatter plot of 22-kHz vocalisation onsets and number of trials. Left and right columns, rats with 20 and 30-s odour presentation, respectively. On the left of each plot is shown the rat identification number and the total number of 22-kHz USVs uttered during the first three and last three trials and during all three periods (pre-odour, odour and post-shock). Yellow shaded area: odour presentation.

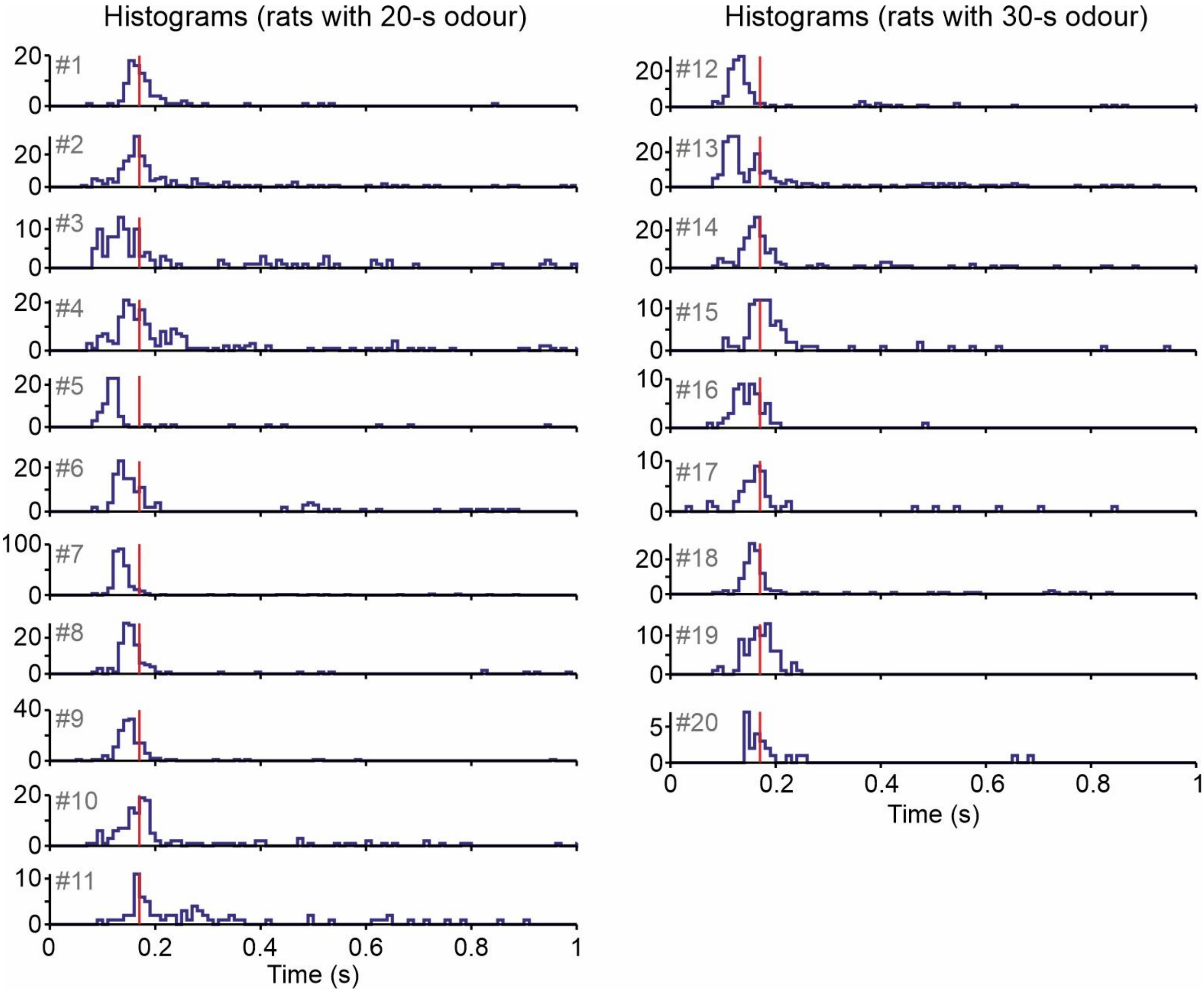


Figure S2. Intercall interval per rat (calculated from offset to onset of the 22-kHz vocalisations). Right and left columns are for the 20 and 30-s odour groups. On the left of each histogram, it is shown the rat identification number. Red line: median of each rat intercall intervals.

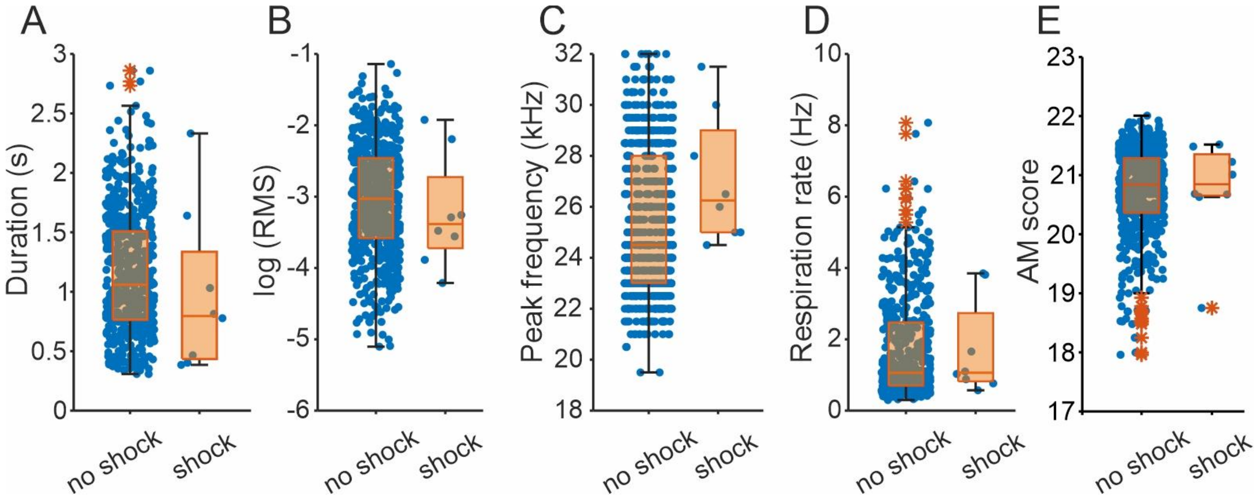


Figure S3. Analyses of 22-kHz USVs and respiration rate during the odour period of fear-conditioning task separating the USVs uttered during the shock and those that were not (during shock, n = 8, no shock, n = 584). A-E) Boxplots (in orange) of the call duration, root mean square (RMS, in natural logarithm), peak frequency, respiration rate and AM score (all variables determined as in the main manuscript). Boxes depict the interquartile range and the horizontal line inside each box indicates the median. The asterisks are outlier values. In blue, each individual value.

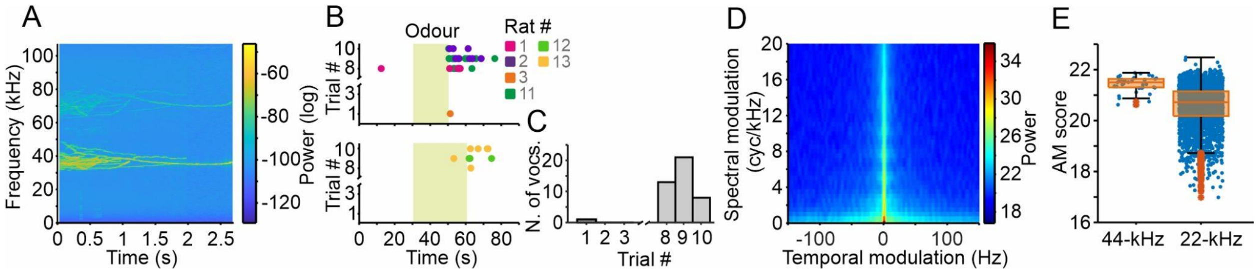


Figure S4. 44-kHz vocalisations. A) Mean spectrogram of the 44-kHz USVs (n=43). B) Scatter plot with the time onset of the vocalisations. Top, 20-s odour presentation; bottom, 30 s; colour-coded for the rat identification number (see legend). C) Histogram of the vocalisations emitted per trial. D) Mean of the modulation power spectra (of those 44-kHz USVs > 300 ms, n=37). E) Amplitude modulation score of 44 (calculated from those used in D) and 22-kHz USVs (for comparison). In orange the boxplot (boxes depict the interquartile range and the horizontal line inside each box indicates the median, asterisks are outlier values), in blue the individual values.

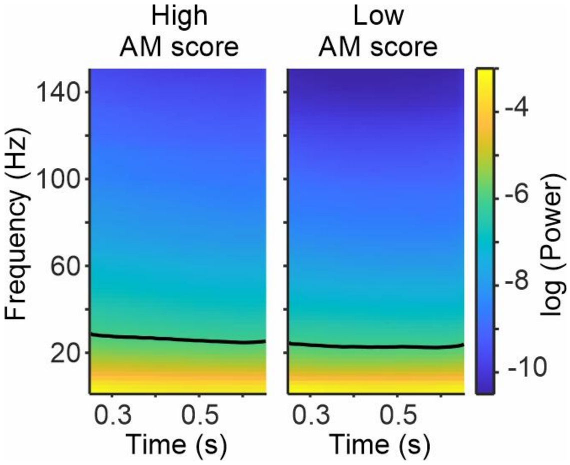


Figure S5. Amplitude modulations remain stable towards the end of the vocalisations. Mean envelope spectrograms of the high and low median split (according to the AM scores) of those vocalisations with duration between 0.9 and 1.5 s (n = 1164; same analysis as in Fig. 5C). Considered here the last 0.9 s.

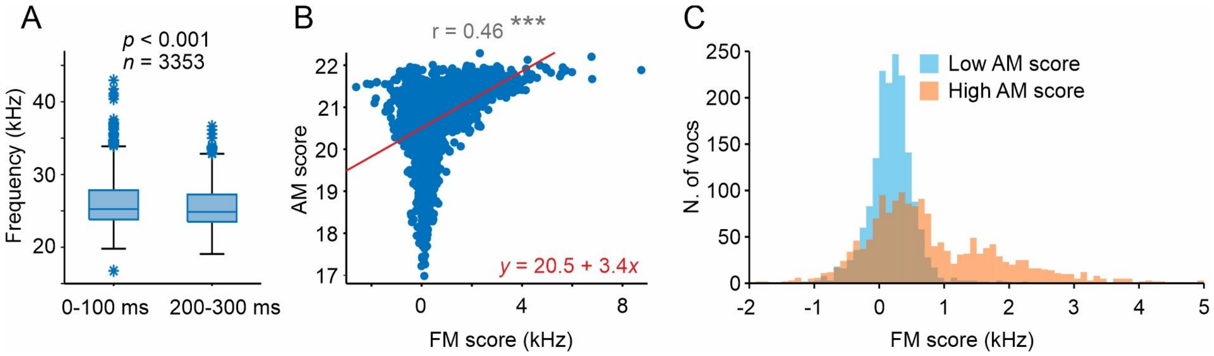


Figure S6. Peak frequency decreases within the first 300 ms. A) Box plot showing the peak frequency within the time windows 0-100 ms and 200-300 ms calculated for all 22-kHz vocalisations (n = 3353, $p < 0.001$, Wilcoxon signed rank test). Boxes depict the interquartile range and the horizontal line indicates the median. B) AM score plotted against the frequency difference (frequency modulation (FM) score), obtained by subtracting "peak frequency at 200-300 ms" from "peak frequency at 0-100 ms". In red, line and equation of a linear regression, bisquare robust fitting. On top, the correlation coefficient ($r$). *** $p < 0.001$. C) Histogram of FM scores. Shown as two distributions for calls with low and high AM scores obtained after median splitting the general AM score distribution. Neither distribution has a median at zero ($p < 0.001$ for both cases, Wilcoxon signed rank test).

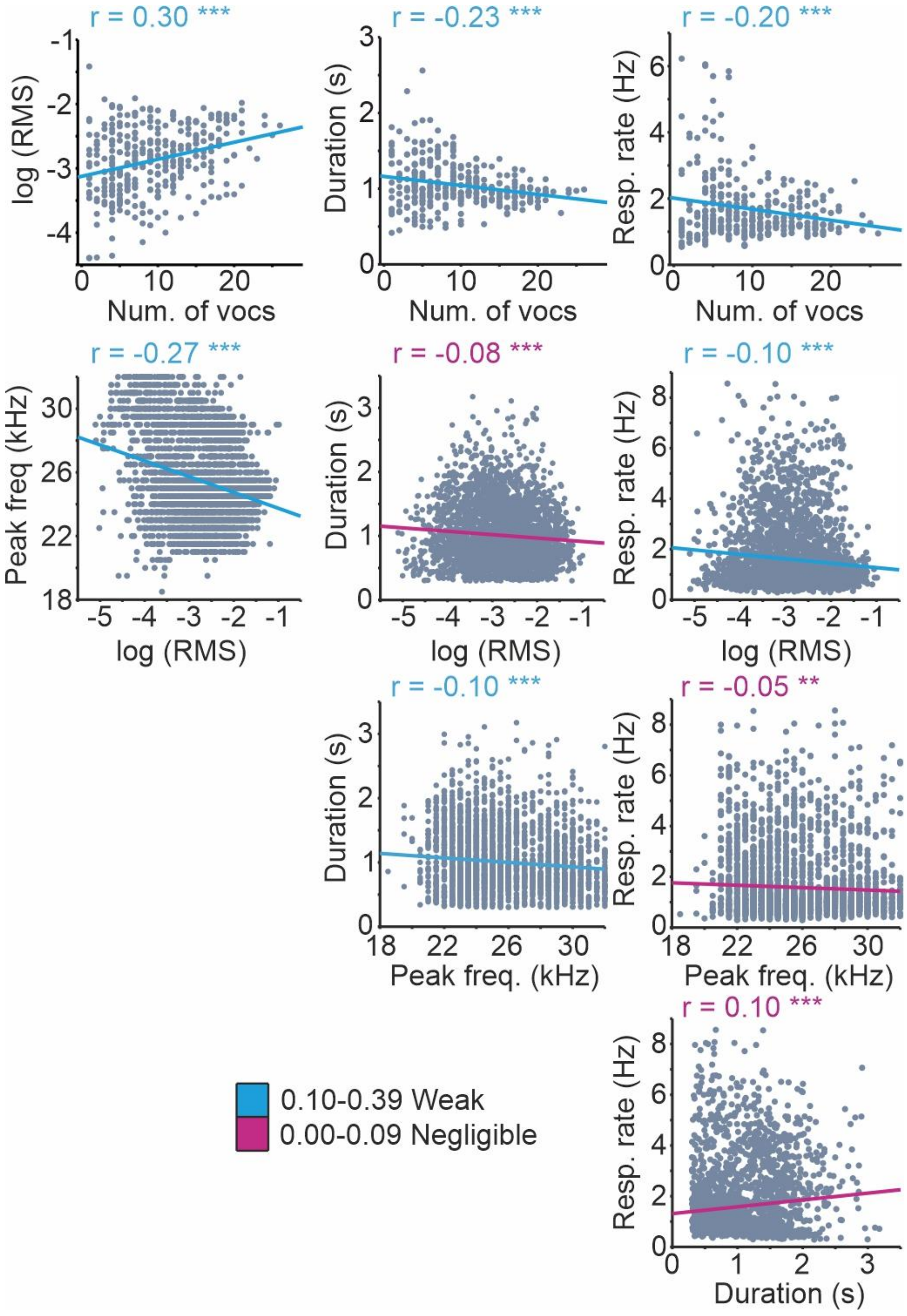


Figure S7. Weak and negligible correlations of 22-kHz vocalisations. Lines depict the fitted linear regression (see colour legend for the interpretation; Schober and Schwarte, 2018). Pearson's correlation coefficient (*r*) shown for each plot. ** $p < 0.01$; *** $p < 0.001$.
